# Engineering Modular Viral Scaffolds for Targeted Bacterial Population Editing

**DOI:** 10.1101/020891

**Authors:** Hiroki Ando, Sebastien Lemire, Diana P. Pires, Timothy K. Lu

**Affiliations:** Department of Electrical Engineering & Computer Science and Department of Biological Engineering, Massachusetts Institute of Technology, 77 Massachusetts Avenue, Cambridge, MA 02139, USA.; MIT Synthetic Biology Center, Massachusetts Institute of Technology, Cambridge MA 02139, USA.; Centre of Biological Engineering, University of Minho, Campus de Gualtar 4710-057, Braga, Portugal.

## Abstract

Bacteria are central to human health and disease, but the tools available for modulating and editing bacterial communities are limited. New technologies for tuning microbial populations would facilitate the targeted manipulation of the human microbiome and treatment of bacterial infections. For example, antibiotics are often broad spectrum in nature and cannot be used to accurately manipulate bacterial communities. Bacteriophages can provide highly specific targeting of bacteria, but relying solely on natural phage isolation strategies to assemble well-defined and uniform phage cocktails that are amenable to engineering can be a time-consuming and labor-intensive process. Here, we present a synthetic-biology strategy to modulate phage host ranges by manipulating phage genomes in *Saccharomyces cerevisiae*. We used this technology to swap multiple modular phage tail components and demonstrated that *Escherichia coli* phage scaffolds can be redirected to target pathogenic *Yersinia* and *Klebsiella* bacteria, and conversely, *Klebsiella* phage scaffolds can be redirected to target *E. coli*. The synthetic phages achieved multiple orders-of-magnitude killing of their new target bacteria and were used to selectively remove specific bacteria from multi-species bacterial communities. We envision that this approach will accelerate the study of phage biology, facilitate the tuning of phage host ranges, and enable new tools for microbiome engineering and the treatment of infectious diseases.

## INTRODUCTION

Bacteriophages (phages) are natural biological nanomachines that have evolved to infect host bacteria with exquisite specificity and efficacy. Phages constitute the most abundant type of biological particles on earth (Hendrix, 2003) and reproduce at the expense of their host bacteria. Thus, phages have been explored as a means of controlling pathogenic bacteria (d’Herelle, 1931), but poor understanding of the molecular relationships between bacteria and their phages can lead to highly variable treatment outcomes (Brussow, 2012). With the rise of drug-resistant bacterial infections and the sharp decline in antibiotic discovery and development (Fischbach and Walsh, 2009), phage therapy is regaining attention after years of declining interest in the Western world (Carlton, 1999). Furthermore, despite the important role that the microbiome plays in regulating human health and disease (Grice and Segre, 2012), strategies for precisely manipulating complex microbial communities are lacking. With their ability to kill or deliver DNA into specific bacteria, phages constitute a promising technology for manipulating microbiota. However, the limited host range of most naturally isolated phages is a major barrier to the development and approval of commercially available phage-based products. Conventional strategies for identifying phages with specific host ranges rely on screening samples from nature. Naturally isolated phages are often very diverse in morphology, genomic content, and life cycles, which poses a challenge for the engineering, manufacturing, and regulatory approval of phages as biotechnologies. For example, phage cocktails have been used to address the limited host range of any single phage (Sulakvelidze et al., 2001). However, the desire to increase coverage by adding more members to a phage cocktail is counterbalanced with the challenge of producing and testing well-defined multi-component mixtures for regulatory approval.

Creating phage-based therapeutics and diagnostics is also limited by the difficulty of engineering phages. Lytic phage DNA does not reside for very long inside of bacteria; this makes it difficult to modify phage genomes during the phage reproductive cycle. Phage genomes are also often too large to be handled *in vitro.* Phage genome engineering is thus classically carried out with allele replacement methods, whereby a piece of the phage genome is cloned into an appropriate bacterial vector and remodeled using classical molecular biology, and the bacterium containing the resulting construct is then infected with the phage, allowing the phage to recombine with the plasmid to acquire the desired mutations. This process is inefficient because many phages degrade resident DNA upon entry, and time-consuming due to the absence of phage selectable markers to expedite screening of output viral populations. Furthermore, there are very large stretches of phage DNA that encode products toxic to bacteria, thus preventing their manipulation within bacterial hosts. Finally, all existing approaches are limited in the number of mutations that can be introduced simultaneously. Multiple rounds of mutations are therefore often required, making the process inefficient. Here, we demonstrate a high-throughput phage-engineering platform that leverages the tools of synthetic biology to overcome these challenges and use this platform to engineer model phages with tunable host ranges.

## RESULTS

### Yeast platform for bacteriophage genome engineering

We used an efficient yeast-based platform (Jaschke et al., 2012; Lu et al., 2013) to create phages with novel host ranges based on common viral scaffolds. Inspired by gap-repair cloning in yeast (Ma et al., 1987) and the pioneering work of Gibson and co-workers (Gibson, 2012; Gibson et al., 2008; Gibson et al., 2009), we captured phage genomes into *Saccharomyces cerevisiae*, thus enabling facile genetic manipulation of modified genomes that can be subsequently re-activated or “rebooted” into functional phages after transformation of genomic DNA into bacteria (Figure 1A). The workflow is split into two parts. In the first part, the entirety of the viral genome to be assembled in yeast is amplified by PCR in such a way that each adjacent fragment has homology over at least 30 bp. The first and last fragments of the phage genome are amplified with primers that carry “arms” that have homology with a yeast artificial chromosome (YAC) fragment, which may be obtained by PCR or any other suitable method. Upon transformation of all viral genome fragments and the YAC into yeast, gap-repair will join each fragment to the adjacent one templated by the homology regions found at the end of each fragment, yielding a full phage genome cloned into a replicative yeast plasmid. Yeast transformants are then enzymatically disrupted in order to extract the YAC-phage DNA, which is then used for transformation into bacterial host cells that can support resumption of the viral life cycle. Plaques, if obtained, are then picked, amplified, and sequenced to verify proper introduction of the desired mutations. If no plaques are obtained, it is still possible to amplify parts of the YAC-phage genome from the yeast clones in order to verify proper DNA assembly, to eliminate the possibility of unwanted mutations, and to help determine potential reasons for the failure of the synthetic phage genome to produce viable offspring.

**Figure 1.**
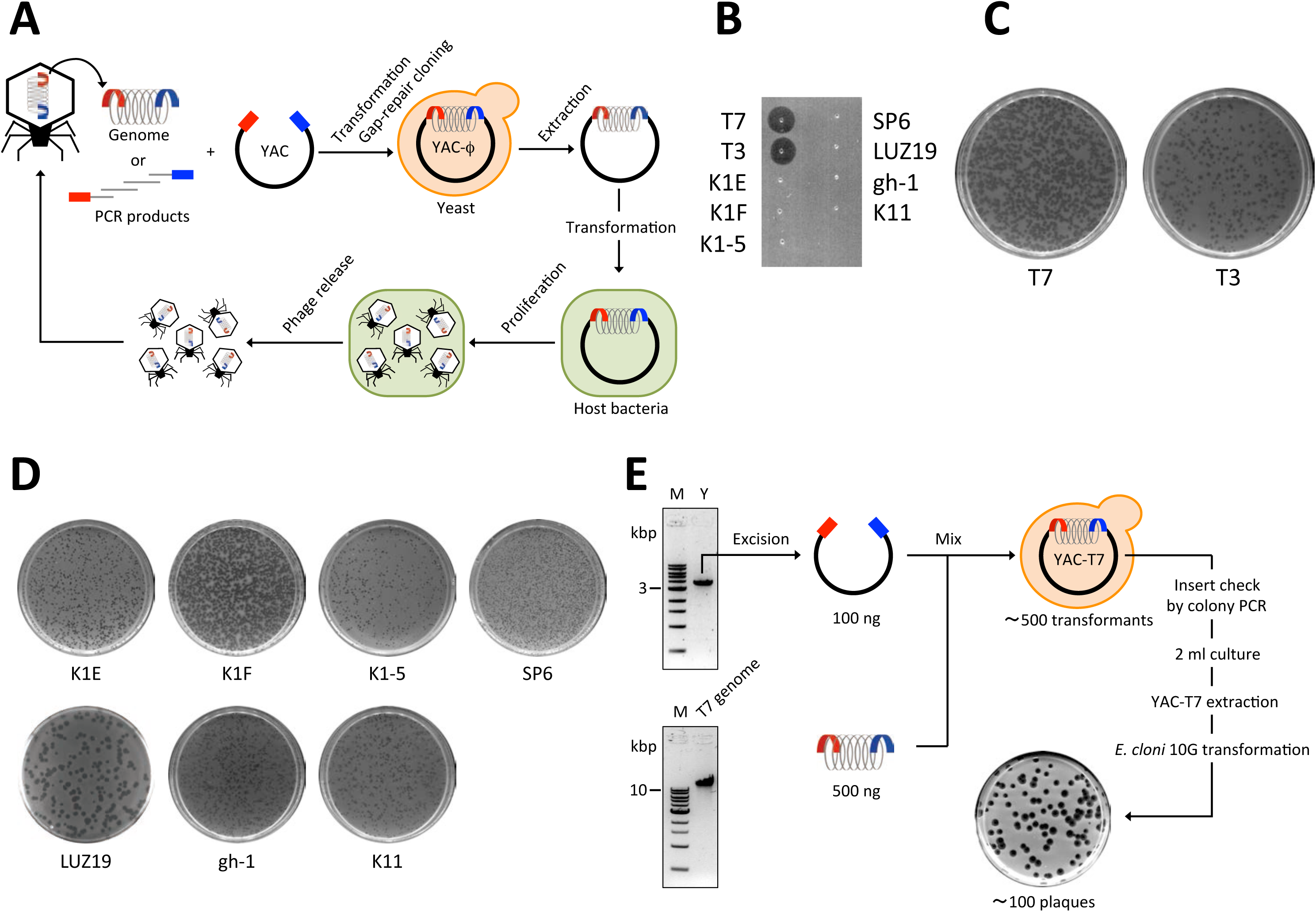
Yeast platform for phage engineering. (**A**) Schematic illustrating the workflow to capture and reboot phages using our yeast platform. An entire phage genome or PCR products spanning an entire phage genome are transformed into yeast cells along with a linearized yeast replicon fragment from the yeast artificial chromosome, pRS415. In yeast, the phage genome is assembled and captured in the YAC by gap-repair cloning. The resulting YAC-phage DNA is extracted and transformed into host bacteria. Active phages are produced from the YAC-phage DNA and generate plaques on a lawn of host bacteria. (**B**) To determine the phage sensitivity of the *E. coli* 10G strain, high-titer phage lysates (>10^9^ PFU/ml) were spotted onto 10G lawns. The 10G strain was sensitive only to infection by T7 and T3 phages. (**C**) Rebooting T7 and T3 phages from purified phage genomes. We electroporated 10 ng of genomic DNA into 10G, mixed chloroform-treated lysates with *E. coli* BL21 and LB soft agar, poured the mixtures onto LB plates, and incubated at 37°C. (**D**) One-time phage propagation assays. Rebooting phages from purified phage genomes via 10G. We electroporated 10-200 ng of phage genomic DNA into 10G and incubated the cells at 37°C for 1 h. We then treated the bacteria with chloroform. After centrifugation, supernatants were mixed with host bacteria and LB soft agar, poured onto LB plates, and incubated at 30 or 37°C. All tested phage genomes, including non-*E. coli* phages, could be rebooted in *E. coli* 10G cells. Host bacteria: IJ1668 K-12 hybrid; K1 capsule for K1E, K1F, and K1-5 phages, IJ612 *Salmonella typhimurium* LT2 for SP6 phage, *Pseudomonas aeruginosa* PAO1 for LUZ19 phage, *Pseudomonas putida* C1S for gh-1 phage, and *Klebsiella* sp. 390 for K11 phage. (**E**) An example of capturing and rebooting a phage through the yeast platform. An excised YAC pRS415 amplicon and the T7 genome were co-transformed in yeast cells. The T7 genome was captured in the YAC by gap-repair cloning. The YAC-T7 DNA was extracted and used for transformation. Progeny phages were produced from YAC-T7 DNA in the *E. coli* 10G strain and generated plaques on *E. coli* BL21.

We targeted phages from the T7-family because their life cycle is largely host independent (Qimron et al., 2010) and there is a relatively large number of family members for which genomic sequences are publicly available. These include T7 (coliphage, 39,937 bp), T3 (coliphage, 38,208 bp), K1E (K1-capsule-specific coliphage, 45,251 bp), K1F (K1-capsule-specific coliphage, 39,704 bp), K1-5 (K1- or K5-capsule-specific coliphage, 44,385 bp), SP6 (*Salmonella* phage, 43,769 bp), LUZ19 (*Pseudomonas* phage, 43,548 bp), gh-1 (*Pseudomonas* phage, 37,359 bp), K11 (*Klebsiella* phage, 41,181 bp), and others. We first sought to confirm that purified phage DNA from various phages could be transformed into bacterial hosts to generate functional phages. We used *E. cloni* 10G® (10G) cells (Durfee et al., 2008) as a one-time phage propagation host. Except for T7 and T3, all phages used in this study (K1E, K1F, K1- 5, SP6, LUZ19, gh-1, and K11) cannot infect 10G (Figure 1B). We extracted the genomes of a diversity of phages from purified phage particles. Each genome was electroporated into 10G directly. After incubation and chloroform treatment, supernatants were mixed with overnight cultures of each natural host bacteria for each phage in soft agar, poured onto agar plates, and incubated, looking for plaque formation (Figure 1C and 1D). We found that all the phages tested could be rebooted from purified DNA into functional phages through one-step propagation in 10G, even if their natural target species was not *E. coli* (Figure. 1D and Table S1). This result indicates that we can use *E. coli* 10G cells as an initial host for rebooting purified genomic DNA into phages that infect diverse bacteria.

### Rebooting bacteriophages from PCR products via the yeast platform

To determine whether phage genomes assembled in yeast remain viable, we first attempted to capture and then reboot T7 (Figure 1E), T3, and LUZ19 phages. We used PCR to amplify the YAC pRS415 and add arms homologous to the ends of the phage genomes. We co-transformed the YAC amplicons with phage genomic DNA into yeast. Confirmed YAC-phage constructs were extracted from yeast and transformed into *E. cloni* 10G. These cells were then chloroform treated and the resulting lysates were assessed for plaque-forming units (pfu) on the natural bacterial hosts of the phages (see Figure 1E for capturing and rebooting T7). All 3 phage genomes yielded yeast clones that could be rebooted to viable phages using this strategy.

Next, we captured and rebooted eight different phages that target *E. coli*, *Salmonella*, *Pseudomonas*, and *Klebsiella* (T7, T3, K1E, K1F, K1-5, SP6, gh-1, and K11) by assembling overlapping 3.8-12 kbp-long PCR products spanning the phage genomes with the linearized YAC in yeast (Figure 2A upper panel illustrates this process with T7 as an example). All eight phages were rebooted from PCR fragments via yeast platform and formed plaques on their natural host bacteria (data not shown). Thus, this approach enables the efficient assembly and instantiation of functional recombinant phages, and allows us to potentially create any desired phage genotype in one step from PCR products.

**Figure 2.**
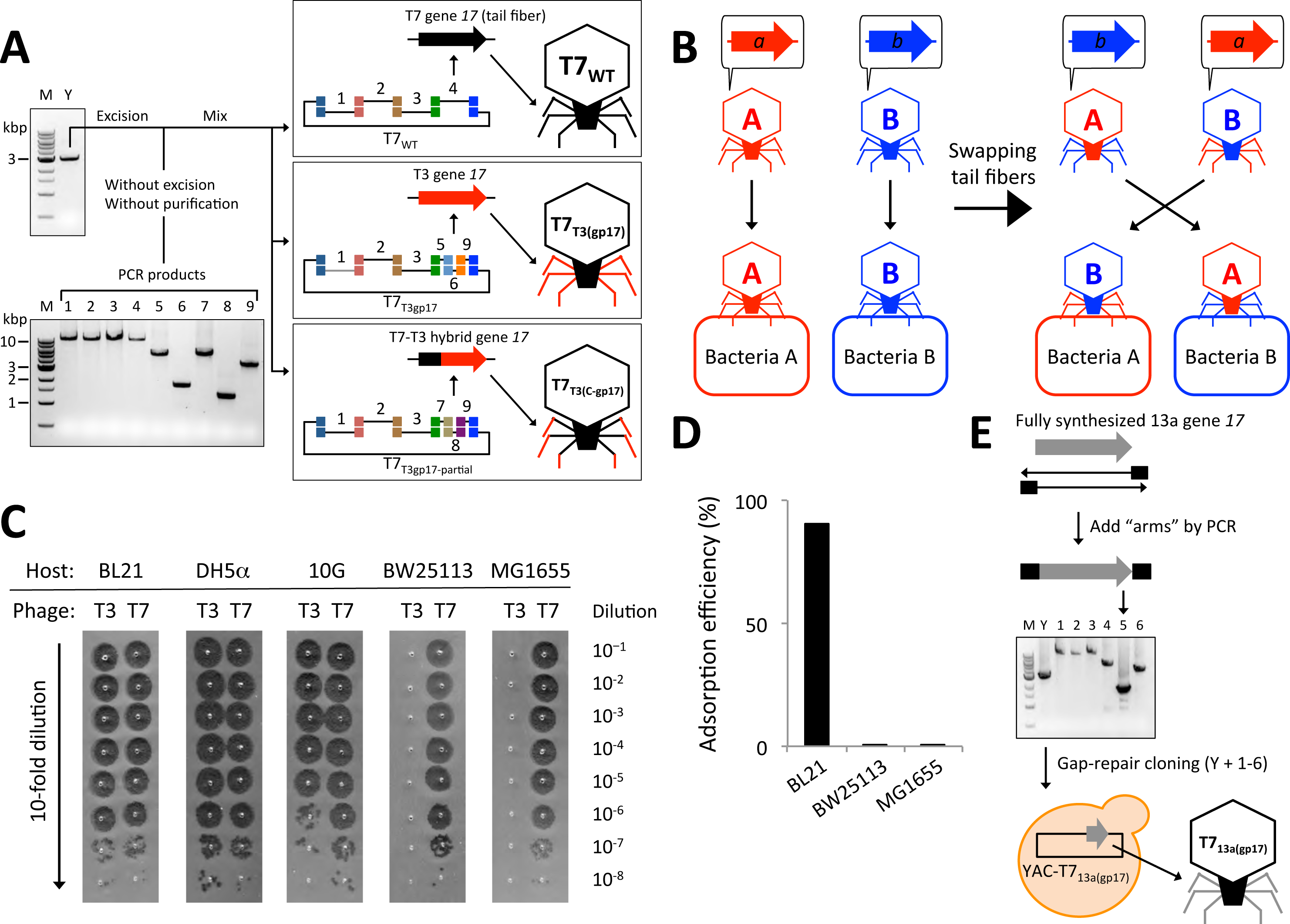
Creation of synthetic phages with engineered host range. (**A**) We prepared multiple PCR fragments encoding the wild-type T7 phage genome (T7_WT_), T7 phage with the entire T3 phage tail fiber (T7_T3(gp17)_), and T7 phage with a hybrid T7-T3 tail fiber (T7_T3(C-gp17)_). All fragments were co-transformed and assembled in yeast along with YAC DNA. M, 1 kb DNA size marker (NEB). Y, YAC amplicon. (**B**) Phage A with its primary host determinant, gene *a*, infects bacteria A, but cannot infect bacteria B. Phage B with its primary host determinant, gene *b*, infects bacteria B, but cannot infect bacteria A. We hypothesized that by swapping these host determinants between phage A and B, engineered phage A with gene *b* and engineered phage B with gene *a* should infect bacteria B and A, respectively. (**C)** Host ranges of T7 and T3 phages. Each bacterial overnight culture and LB soft agar were mixed, and poured onto LB plates. 2.5 μL of 10-fold serially diluted T7 and T3 phages were spotted onto the bacterial lawns and incubated at 37°C. T3 phage did not plaque efficiently on *E. coli* BW25113 and MG1655, whereas T7 phage plaqued efficiently on all tested *E. coli* strains. (**D**) Adsorption assay. Bacteria and T3 phage were mixed at an MOI ∼0.5 and incubated for 10 min. Growth of adsorbed progeny was stopped by the addition of chloroform. After centrifugation, supernatants were serially diluted and mixed with *E. coli* BL21 and LB soft agar, and poured onto LB plates. After incubation at 37°C, phage plaques were counted, and adsorption efficiencies were calculated. The data are presented as the mean of three independent experiments, and the error bars represent the SEM. Small error bars are obscured by bar charts. (**E**) Creation of synthetic T7 phage with phage 13a tail fiber (encoded by gene *17*). We synthesized 13a’s gene *17* and assembled it with the rest of the T7 genome via overlapping PCR products in yeast. The YAC-phage DNA was extracted and used for transformation. M, 1 kb DNA size marker (NEB). Y, YAC amplicon.

### Swapping tail fibers enables modulation of phage host range

To engineer phages with tunable host ranges, we first selected two model phages, T7 and T3, which are obligate lytic phages originally isolated as a member of the seven “Type” phages that grow on *E. coli* B (Demerec and Fano, 1945). T7 and T3 have linear genomes that share high homology with each other, in which the primary host determinant is the product of gene *17* (gp17), the tail fiber (Dunn and Studier, 1983; Pajunen et al., 2002). Alterations in the gp17 sequence have been linked to the recognition of different host receptors and shifting host ranges (Molineux, 2006). Thus, we hypothesized that exchanging gene *17* or fragments of gene *17* between T7, T3, and their relatives could be used to tune their host specificities (Figure 2B). This is supported by previous data on naturally occurring hybrids between T7 and T3 whose host range was mostly dictated by which gene *17* they harbored (Lin et al., 2012).

We first examined the host range of T7 and T3 phage on a range of hosts to determine bacterial panels that were differentially targeted by the two phages. T3 is described as incapable of targeting many common laboratory *E. coli* K-12 strains (Molineux, 2006), so we performed plaque formation assays with four K-12 strains and a B strain (BL21) as a control. As shown in Figure 2C, T7 plaqued efficiently on all strains, while T3 did not produce plaques on BW25113 and MG1655 at a detectable frequency (Efficiency Of Plating (EOP) below 10^-9^). T3 exhibited ∼4 orders-of-magnitude reductions in adsorption efficiency on BW25113 and MG1655 compared with the permissive BL21 strain (Figure 2D). These results indicate that we can differentiate between T7 and T3 using BW25113 or MG1655, which are only susceptible to T7.

The gp17 tail fibers of T7 and T3 can be split in two domains. The N-terminal 149 residues are thought to be necessary for the tail fiber to bind to the rest of the capsid while the remaining C-terminal region forms a kinked shaft and harbors the recognition domain for host receptors at its tip (Steven et al., 1988). The N-terminal regions of T7 and T3 share 99% identity at the protein level, while the C-termini exhibit 83% identity, with the last 104 amino acids of the T3 protein showing only 62% of identity to the corresponding 99 amino acids of the T7 protein. Therefore, we hypothesized that swapping the C-terminal domain between the two viruses would result in exchanging the host ranges. We constructed synthetic phages, based on either the T7 or T3 viral chassis, which carried engineered gene *17* alleles composed of fragments from the other phage. Specifically, we created six synthetic phages: T7 phage with the wild-type T7 tail fiber (T7_WT_), T7 phage with 410 amino acids of the C-terminal region of the T3 tail fiber (T7_T3(C-gp17)_), T7 phage with the entire T3 tail fiber (T7_T3(gp17)_), T3 phage with the wild-type T3 tail fiber (T3_WT_), T3 phage with 405 amino acids of the C-terminal region of the T7 tail fiber (T3_T7(C-gp17)_), and T3 phage with the entire T7 tail fiber (T3_T7(gp17)_). T7_WT_ and T7 phages are the same at the genetic level; however, T7_WT_ phage was created by capturing the T7 genome in yeast and then rebooting this phage genome in bacteria and served as a control for the faithfulness of the reconstruction process, whereas T7 was obtained from ATCC. The same applies to T3_WT_ and T3. Each phage was assembled in yeast via six PCR fragments and was rebooted via transformation into *E. coli* 10G (example schematics in Figure 2A). No unexpected mutations were found in the heterologous gp17 regions of the rebooted phages.

To examine the host specificities of our six engineered phages, we performed plaque formation assays on a range of *E. coli, Klebsiella*, and *Yersinia pseudotuberculosis* (*Y. ptb*) strains (Figure 3). T3_T7(C-gp17)_ and T3_T7(gp17)_ plaqued on *E. coli* BW25113 and *E. coli* MG1655 at a similar EOP as T7 and T7_WT_, while T3, T3_WT,_ T7_T3(C-gp17)_, and T7_T3(gp17)_ had >10^5^-fold-reduced EOPs on these strains. In addition, T3, T7_T3(C-gp17)_, and T7_T3(gp17)_ plaqued on *E. coli* ECOR16 while T7, T3_T7(C-gp17)_, and T3_T7(gp17)_ did not. In addition, we also synthesized a codon-optimized version of the tail fiber of the T7-like enterobacteriophage 13a and created synthetic T7 phages containing the entire 13a tail fiber (T7_13a(gp17)_) or the C-terminal region of the 13a tail fiber (T7_13a(C-gp17)_) (Figure 2E, Table S2). Although T7 and T7_WT_ did not plaque on *E. coli* ECOR16, both T7_13a(gp17)_ and T7_13a(C-gp17)_ were able to do so efficiently (Figure 3). These results demonstrate that the C-terminal region of gp17 is a major host range determinant and that new host ranges can be conferred onto T7-like phage scaffolds by engineering tail fibers. Interestingly, T7_13a(C-gp17)_ efficiently infected *E. coli* BW25113 and MG1655, similar to T7 and T7_WT_, but T7_13a(gp17)_ did not, which suggests that the N-terminus of the phage 13a tail fiber can also alter infectivity of the virus although the mechanism is still to be investigated. A second example of this phenomenon can be found between T7_T3(C-gp17)_ (lane 3, Figure 3) and T7_T3(gp17)_ (lane 4, Figure 3). While the former phage infected *Y. ptb* YPIII (albeit with a low EOP), the latter phage as well as wild-type T3 did not.

**Figure 3.**
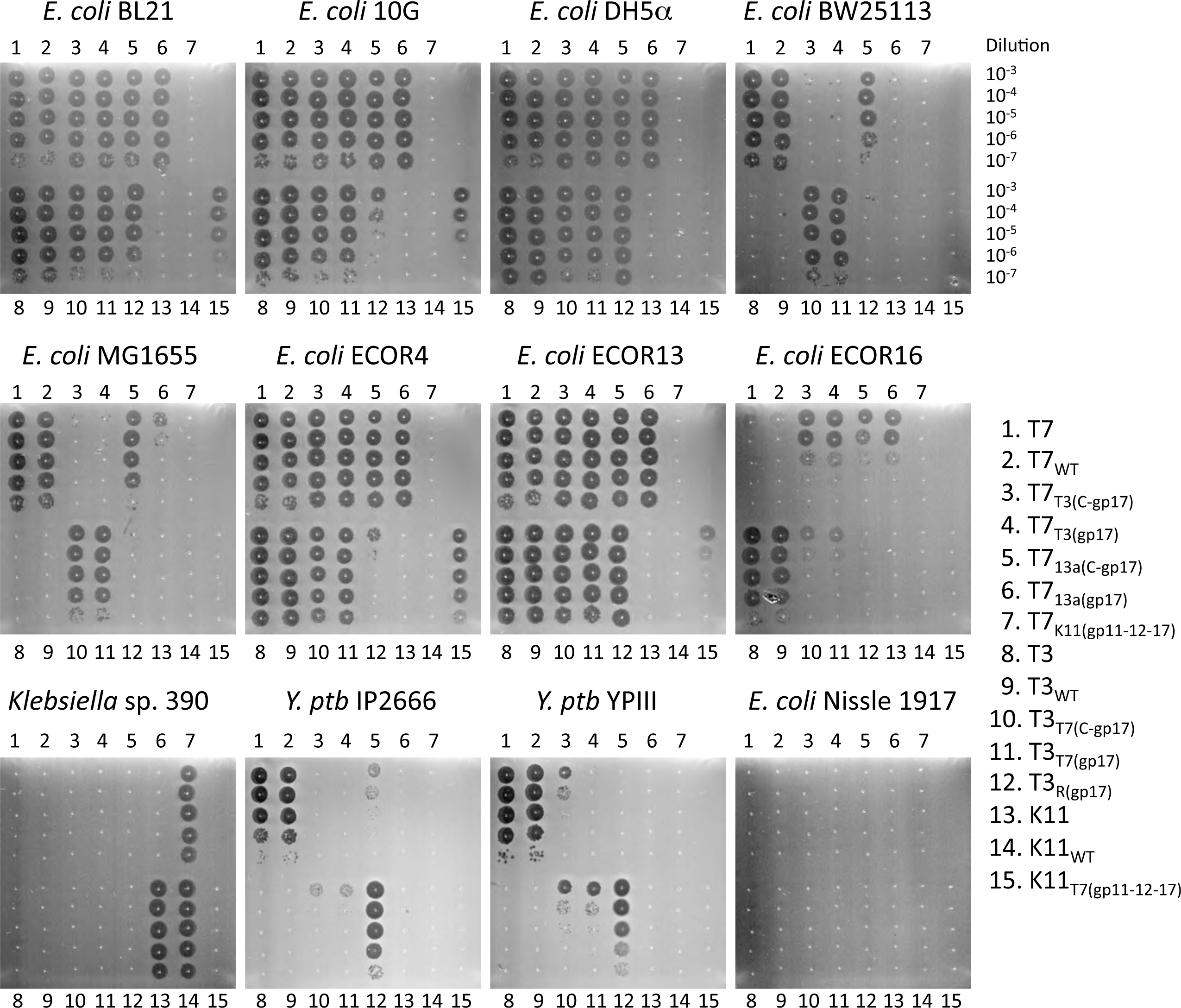
Plaque formation assays with natural, reconstructed wild-type, and synthetic phages. Bacterial lawns were spotted with 2.5 μL of 10-fold serially diluted phages and incubated at 30 or 37°C. Synthetic phages showed tail-fiber- or tail-component-dependent host ranges and the ability to cross between species.

### Coliphage T3 with a *Yersinia* phage tail fiber infects both *E. coli* and *Y. pseudotuberculosis*

We further demonstrated that gene swapping between phages could overcome species barriers by designing synthetic phage based on T7 or T3 scaffolds that can infect bacteria other than *E. coli*. We started with coliphage T3 and *Yersinia* phage R (38,284 bp), since their gp17’s share 99.5% identity at the protein level and differ by only 3 nucleotides in gene *17*. We hypothesized that these differences could be responsible for their divergent host ranges. Indeed, we were unable to detect productive T3 infection of *Y. ptb* strains IP2666 and YPIII, which are known hosts for phage R (Rashid et al., 2012). Because we did not have access to phage R, we introduced the desired mutations in T3 gene *17* by PCR so that it would encode the same tail fiber as phage R (Figure 4A). The mutated gene *17* was then swapped into the genome of T3. Synthetic T3 phage with the R tail fiber (T3_R(gp17)_) was able to infect *Y. ptb* IP2666 and YPIII. Interestingly, T3_R(gp17)_ maintained the capacity to infect *E. coli* BL21 (Figure 4B), demonstrating that the introduced mutations conferred a host range expansion and not just a host range shift. In addition to plaquing assays, we further characterized the ability of T3_WT_ versus T3_R(gp17)_ to kill *Y. ptb* IP2666 over time. After 1.5 h of treatment, T3_R(gp17)_ killed 99.999% of IP2666 while T3 had no effect on the bacteria (Figure 4C).

**Figure 4.**
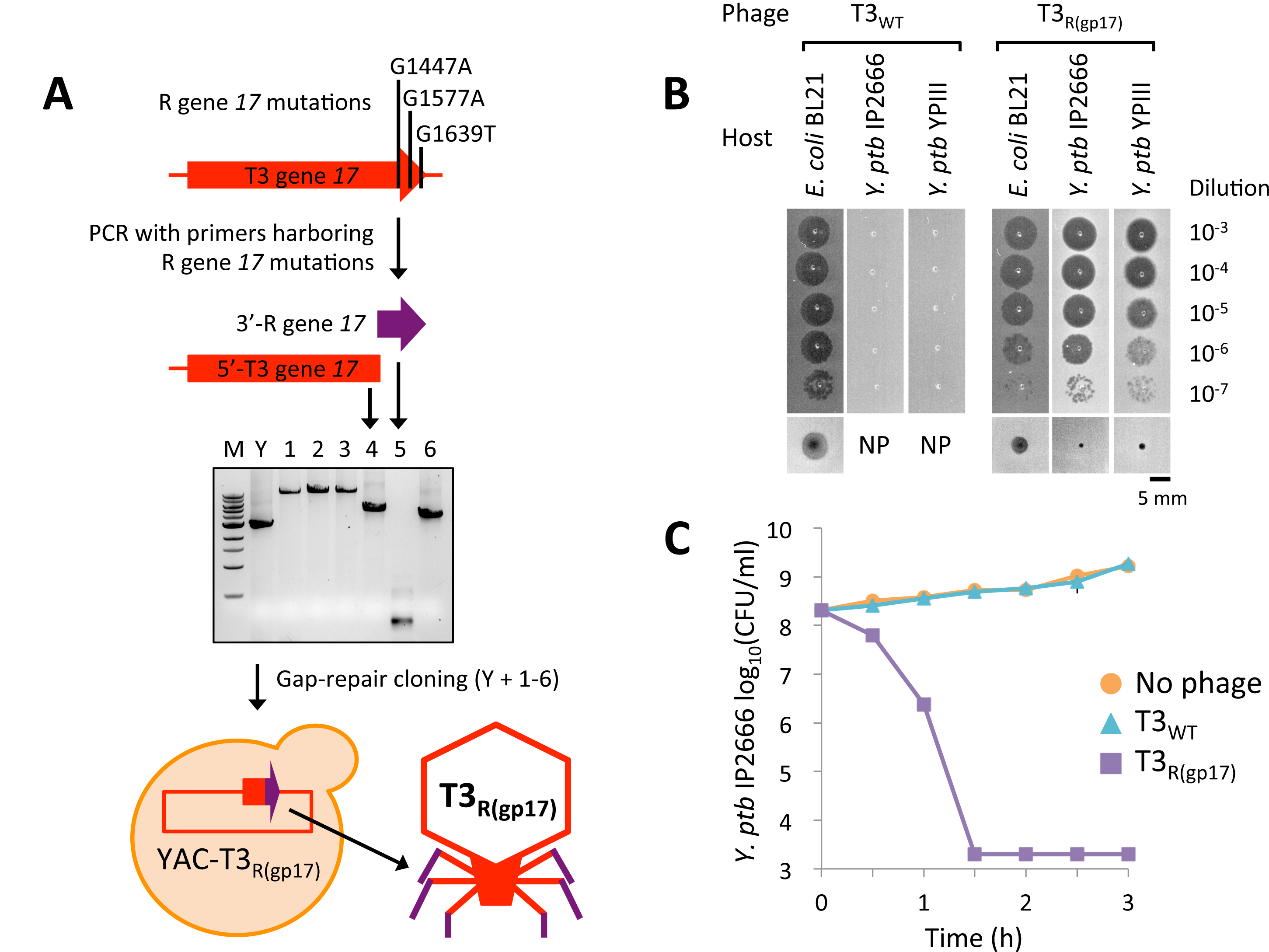
Creation of synthetic T3 phage with *Yersinia* phage R tail fiber. (**A**) We introduced mutations in T3 gene *17* PCR to convert it into phage R gene *17* and assembled the resulting product with the rest of the T3 genome and YAC DNA in yeast. The YAC-phage DNA was extracted and used for transformation into *E. coli*. M, 1 kb DNA size marker (NEB). Y, YAC amplicon. (**B**) Plaquing assay with T3_WT_ and T3_R(gp17)_ on *E. coli* BL21, *Y. ptb* IP2666, and *Y. ptb* YPIII demonstrates that T3_R(gp17)_ has the ability to infect both *E. coli* and *Y. ptb*. Ten-fold serial dilutions of phage lysates were spotted on bacterial lawns and incubated for 4 h at 37°C for *E. coli* BL21 or 24 h at 30°C for *Y. ptb* strains. These pictures were cut out from Figure 3. Bottom panels show images of individual plaques. NP, no plaque. (**C**) Killing curves of *Y. ptb* IP2666 treated with T3_R(gp17)_. ∼10^8^ CFU/ml bacteria and ∼10^7^ PFU/ml phage were used (MOI ∼0.1). The data are presented as the mean of three independent experiments, and the error bars represent the SEM. Small error bars are obscured by symbols. The detection limit was 2.0 × 10^3^ CFU/ml.

### Redirection of host range between coliphage and *Klebsiella* phage by swapping whole tail components

We further explored our ability to overcome species barriers by engineering phages with lower similarity with one another. K11 is a *Klebsiella* phage that belongs to the T7-like family (Dietz et al., 1990). K11 has a similarly sized genome to T7 and shares gene synteny with T7. The average homology between K11 and T7 is 59% among the genes that have homologs between the two viruses. For comparison, T7 and T3 share 72% identity at the genomic level between homologous genes. While T7 is a coliphage and does not infect *Klebsiella*, K11 infects *Klebsiella*, such as *Klebsiella* sp. 390, but not *E. coli* (Figure 3) (Bessler et al., 1973). Their respective host range determinants, gp17, are very different and do not share any homology outside of the N-terminal 150 amino acids, which is only 47% identical between the two proteins. Specifically, the T7 gp17 encodes tail fibers while K11 gp17, which is 322 amino acids longer than the T7 gp17, directs the synthesis of a tail spike, an enzymatic host range determinant that actively breaks down the capsule of *Klebsiella* to allow K11 phage to gain access to unknown secondary receptors located beneath the capsule (Bessler et al., 1973).

To create a T7 phage with a K11 tail fiber and a K11 phage with a T7 tail fiber, we first swapped the entire gene *17*, but this yielded no viable phages. We then tried to construct composite tail fibers composed of gene *17* fragments from both phages hybridized at various points along the length of the gene, but this was also unsuccessful at generating functional synthetic phages. We speculated that one possible reason for these failures could be that the K11 genome cannot create productive phages within *E. coli* 10G. However, the natural K11 genome produced functional virions when it was electroporated into 10G cells, which were subsequently lysed via chloroform and plated onto a suitable host (Figure 1D and Table S1).

Alternatively, the gene *17* product from K11 may require a function or factor that is absent from T7. Cuervo *et al.* reported that the tail of T7 phage, which assembles independently of the head, is assembled from a dodecamer of gp11 (the adaptor) and an hexamer of gp12 (the nozzle) (Figure 5A) onto which 6 trimers of gp17 attach (Cuervo et al., 2013). T7’s six tail fibers attach at the interface between the adaptor and nozzle, thus making contacts with both proteins. The adaptor ring is responsible for the attachment of the preformed tail to the prohead via interactions with the portal composed of 12 subunits of gp8. The homology between the gp8 of T7 and K11 (80% identity at the amino acid level) is much higher than the homology between the gp11 and gp12 proteins of T7 and K11 (60 and 61% identity, respectively), which led us to suspect that replacing all three tail genes of T7 with their K11 equivalents (gp11, gp12, and gp17) could be necessary to create functional virions (Figure 5B). Indeed, both T7 with K11 tail components (T7_K11(gp11-12-17)_) and K11 with T7 tail components (K11_T7(gp11-12-17)_) were successfully engineered into functional phages that exhibited tail-dependent host ranges. Specifically, T7_K11(gp11-12-17)_ infected *Klebsiella* sp. 390 and did not target *E. coli*, while K11_T7(gp11-12-17)_ infected *E. coli*, but did not plaque on *Klebsiella* (Figure 5C and Figure 3). The yeast-based phage engineering platform enabled the facile construction of these phages via one-step genome construction even though gene *11* and *12* are physically separated from gene *17*, a feat that no other phage engineering method can currently achieve. To further validate the ability of synthetic T7_K11(gp11-12-17)_ to target *Klebsiella*, we performed a time-course experiment that showed that T7_K11(gp11-12-17)_ killed 99.955% of *Klebsiella* sp. 390 after 1 hour of treatment (Figure 5D), but was about 100-fold less effective than K11_WT_ (Figure 5C and Figure S1).

**Figure 5.**
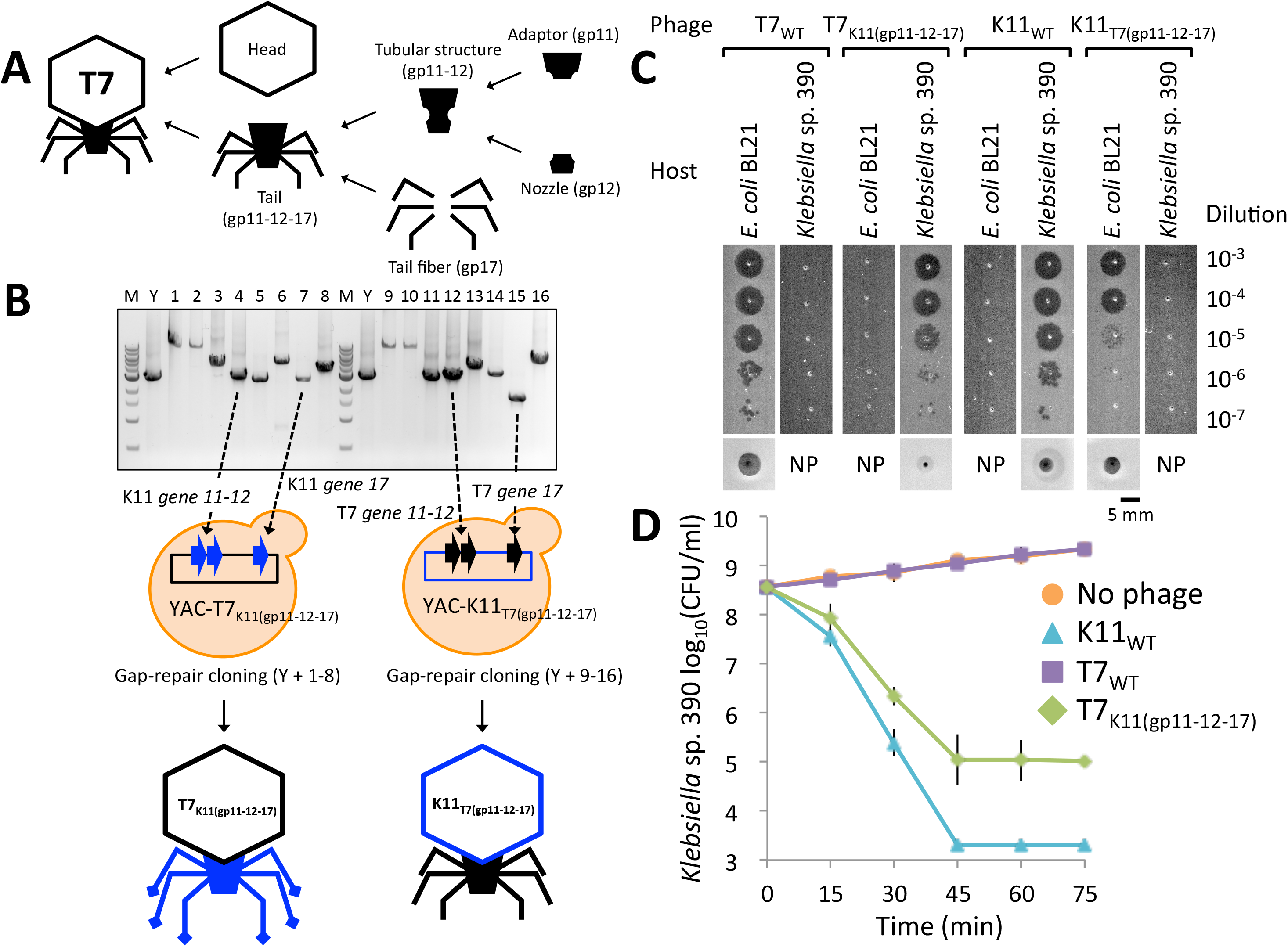
Creation of synthetic T7 phage with *Klebsiella* phage K11 tail components as well as K11 phage with T7 tail components. (**A**) The tail complex of T7 phage is composed of two components, a tubular structure and tail fibers. The tubular structure consists of an upper dodecameric ring made of adaptor protein gp11 and a pyramidal hexameric complex of the nozzle protein gp12. The tail fiber protein gp17 interacts with the interface between gp11 and gp12 (Cuervo et al., 2013). (**B**) Schematics illustrating the construction of synthetic hybrids between phages T7 and K11. Whole genomes were amplified as overlapping PCR amplicons as shown on the gel. Appropriate fragments were co-transformed and assembled in yeast. YAC- phage genomes were extracted and used for transformation. We swapped K11 genes *11*, *12* and *17* into T7 to create T7_K11(gp11-12-17)_ and T7 genes *11, 12* and *17* into K11 to create K11_T7(gp11-12-17)_. M, 1 kb DNA size marker (NEB). Y, YAC amplicon. (**C**) Plaquing of T7_K11(gp11-12-17)_ and K11_T7(gp11-12-17)_ on *E. coli* BL21 and *Klebsiella* sp. 390. Ten-fold serial dilutions of phage lysates were spotted on bacterial lawns and incubated for 4 h at 37°C. These pictures were cut from Figure 3. Bottom panels show images of individual plaques. NP, no plaque. (**D**) Killing curves of *Klebsiella* sp. 390 treated with T7_K11(gp11-12-17)_. ∼10^8^ CFU/ml bacteria and ∼10^7^ PFU/ml phage were used (MOI ∼0.1). The data are presented as the mean of three independent experiments, and the error bars represent the SEM. Small error bars are obscured by symbols. The detection limit was 2.0 × 10^3^ CFU/ml.

### Synthetic phage cocktails efficiently remove target bacteria from mixed bacterial populations

Our results demonstrate that common phage scaffolds can be retargeted against new bacteria hosts by engineering single or multiple tail components. This capability enables the construction of defined phage cocktails that only differ in their host-range determinants and can be used to edit the composition of microbial consortia and/or treat bacterial infections. To demonstrate microbial population editing, we used our synthetic phages to specifically remove targeted host bacteria from a mixed population containing the probiotic *E. coli* strain Nissle 1917, *Klebsiella* sp. 390, and *Y. ptb* IP2666. The amount of each bacterial member in this mixed population was quantified using their differing sensitivities to chemical antimicrobials (Figure S2). *Klebsiella* sp. 390 is naturally resistant to 25 μg/ml carbenicillin while *Y. ptb* IP2666 is naturally resistant to 1 μg/ml triclosan (Figure S2A). *E. coli* Nissle 1917 however is sensitive to both. These concentrations of antimicrobials completely killed susceptible strains but did not visibly affect the growth of resistant strains (Figure S2B). After 1 h treatment of the multi-species population with T7_K11(gp11-12-17)_ or T3_R(gp17)_, >99.9% or >98% of their target bacteria, *Klebsiella* sp. 390 or *Y. ptb* IP2666, respectively, were removed without affecting the remaining bacterial species (Figure 6 and Table S3). Furthermore, a phage cocktail consisting of two phages with the same chassis but different host ranges, T7_WT_ and T7_K11(gp11-12-17)_, resulted in >99% killing of *Klebsiella* sp. 390 and >99.9% killing of *Y. ptb* IP2666 after 1 hour, thus enriching for probiotic *E. coli* Nissle 1917 (Figure 6 and Table S3). These results demonstrate the high efficiency and selectivity of our engineered phages in microbial consortia, and the potential for combining well-defined phage cocktails with probiotics.

**Figure 6.**
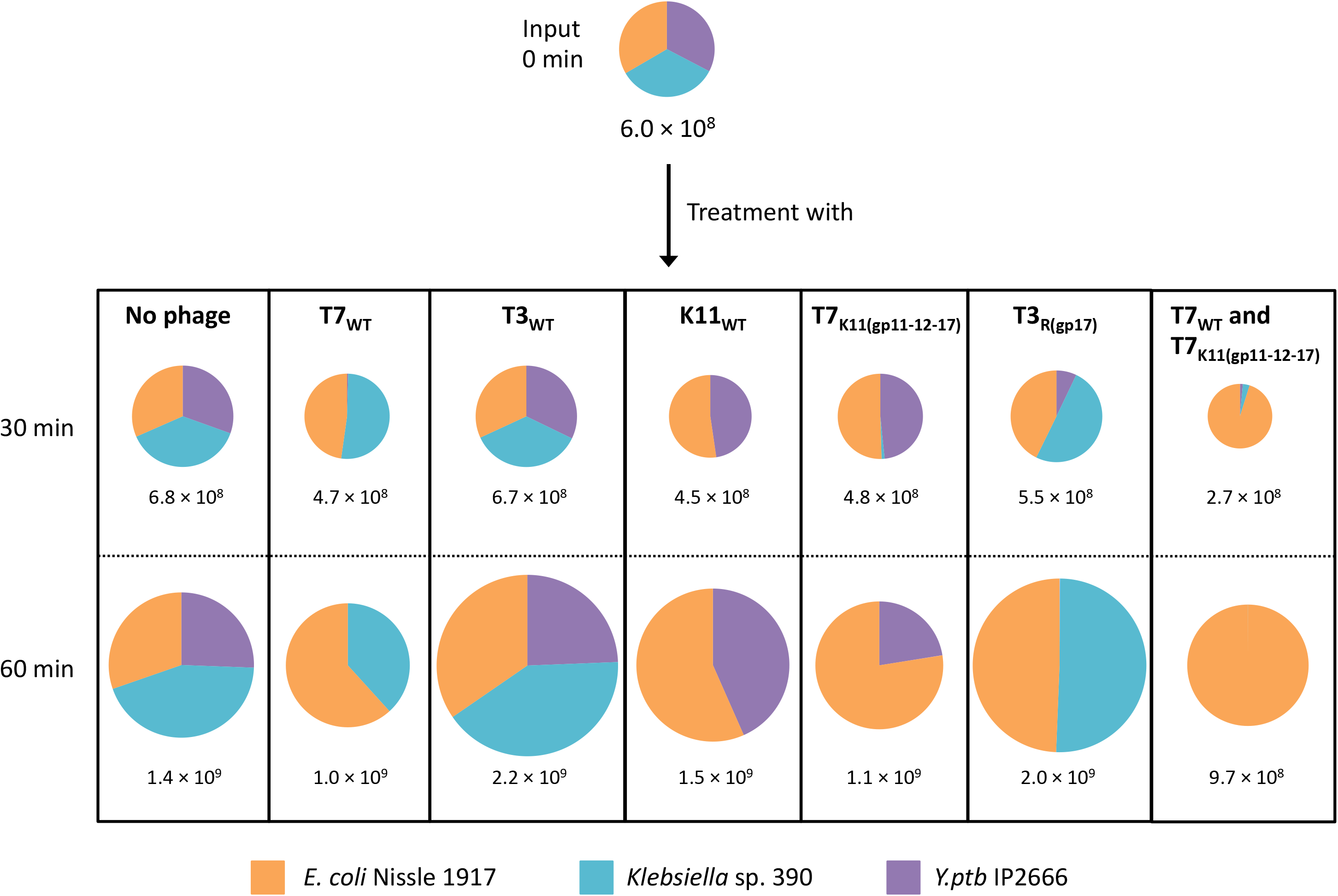
Microbiome editing assay. A synthetic microbial community composed of *E. coli* Nissle 1917, *Klebsiella* sp. 390, and *Y. ptb* IP2666 was treated with various individual synthetic phages and a pairwise combination of phages. After adding ∼10^7^ PFU/ml of each phage, the resulting samples were incubated at 30°C with shaking for 1 h. At each time point, bacteria were collected, washed in saline, serially diluted, and plated onto selective plates for viable cell counts after a 24 h incubation at 30°C. The data are presented as the mean of three independent experiments and the total numbers of cells (CFU/ml) are shown. The sizes of the pie charts reflect the total number of cells. Note that the chart does not allow the display of fractions smaller than ∼1%. The detailed data and the SEM are shown in Table S3. The detection limit was 2.0 × 10^3^ CFU/ml.

## DISCUSSION

In this study, we utilized an efficient yet simple yeast-based platform for phage engineering to modulate phage host ranges for several members of the T7 phage family. Traditional phage engineering strategies, such as *in vitro* manipulation, allele-exchange within bacterial hosts, and phage crossing via co-infection of bacteria (Beier et al., 1977; Garcia et al., 2003; Lin et al., 2011) have been used to modulate phage host range (Tetart et al., 1998; Trojet et al., 2011; Yoichi et al., 2005), but these strategies are inefficient and unable to achieve multiple genetic modifications in a single step. Screening for a desired mutation after classical crossing or recombination experiments can require PCR, restriction digestion, or plaque hybridization on hundreds of individual plaques, which are all costly and time-consuming methods. Conversely, our strategy rarely requires the screening of more than a few yeast clones. Specifically, we found that at >25% of our yeast clones contained properly assembled phage genomes (composed of up to 11 DNA fragments) that could be used to generate functional plaques after transformation into bacteria. Previously, a scheme for engineering phage T4 through electroporation of PCR products was devised (Pouillot et al., 2010), but it is based on a particular feature of the genetic regulation of T4 and cannot easily be applied to other phage families. Recently, the 5.4 kb filamentous coliphage ϕX174 was assembled in yeast in order to stably store the genome and aid in phage refactoring (Jaschke et al., 2012). In this approach, the majority of the genome assembly was performed *in vitro* and the YAC cloning was mostly used to store the resulting genome, whereas the majority of the genome engineering in our approach stems from the actual gap-repair cloning process in yeast. In addition, the phages we have cloned using this method are in the 38-45 kbp range and we have indications that it can also be used for much larger phage genomes (e.g., up to 100 kbp, data not shown).

More recently, type I-E CRISPR-Cas counter-selection has been shown to be a useful tool to edit the genome of phage T7 (Kiro et al., 2014). The *S. pyogenes* CRISPR-Cas9 system was also shown to be functional on the heavily modified genomes of a few members of the T-even family, suggesting that it could be used to modify their genomes, although the authors did not report any such attempts (Yaung et al., 2014). We have successfully used the *Streptococcus pyogenes* CRISPR-Cas9 system to select for mutants in phage T7 but with variable efficiencies (data not shown). Thus, CRISPR-Cas systems can help to overcome some challenges associated with engineering phage genomes in bacterial hosts for therapeutic applications (Bikard et al., 2014; Citorik et al., 2014; Goldberg et al., 2014; Kiro et al., 2014). In contrast, leveraging yeast to modify phages enables the decoupling of phage genome engineering from phage fitness and viability, obviates the need for selective or screenable markers in phage genomes, reduces the risks of phage contamination during the engineering process, and permits facile one-step genetic manipulations. For example, the ability to simultaneously engineer multiple loci in a phage genome was crucial for constructing K11_T7(gp11-12-17)_ and T7_K11(gp11-12-17)_. However, a challenge of yeast-based phage engineering (which is shared by *in vitro* strategies) is the need to reboot modified phage genomes into functional phages. Here, we used high-efficiency DNA transformation to deliver phage DNA into bacterial hosts, but future work may be facilitated by *in vitro* transcription-translation systems capable of supporting functional phage synthesis (Shin et al., 2012).

In summary, we demonstrated that synthetic phages based on common viral scaffolds can be designed to target a range of different bacterial hosts. Furthermore, we showed that a cocktail containing multiple engineered phages could effectively remove select bacterial targets in mixed microbial populations. We anticipate that the systematic and high-throughput engineering of viral genomes will enable new applications and enhanced understanding of bacterial viruses. For example, the engineering of common viral scaffolds could help simplify the discovery and manufacturing of novel bacteriophages and reduce the regulatory burden required for the use of phage cocktails as human therapeutics. Furthermore, abundant phage sequences contained within metagenomic databases could be synthesized and booted into functional phage particles for study and use. Finally, the systematic deconstruction and manipulation of these viral nanomachines will enable a greater understanding of phage biology and may provide insights that are useful for bioinspired nanotechnologies.

## EXPERIMENTAL PROCEDURES

**Strains, vector, and primers.** Phages T7 (ATCC BAA-1025-B2, NC_001604) and T3 (ATCC 110303-B3, AJ318471) were laboratory stocks. Phages K1E (NC_007637), K1F (NC_007456), K1-5 (NC_008152), SP6 (NC_004831), and K11 (EU734173) were kindly provided by Ian Molineux (The University of Texas at Austin). Phage LUZ19 (NC_010326) was kindly provided by Rob Lavigne (KU Leuven). Phage gh-1 (ATCC 12633-B1, NC_004665) was obtained from ATCC. Synthetic phages are listed in Table S4. *Saccharomyces cerevisiae* BY4741 (*MAT*a *his3*Δ*1 leu2*Δ*0 met15*Δ*0 ura3*Δ*0*) was obtained from Thermo Scientific. *Escherichia coli* strains BL21 [B, F^-^ *ompT hsdS*_B_ (r_B_^−^ m_B_^−^) *gal dcm*], DH5α [K–12, F^-^ λ^-^ Φ80d *lacZ*ΔM15 Δ(*lacZYA*- *argF*)U169 *deoR recA*1 *endA*1 *hsdR*17 (r_K_^−^ r_K_^+^) *phoA supE*44 *thi*-1 *gyrA*96 *relA*1], BW25113 [K-12, F^-^ λ^-^ Δ(*araD*-*araB*)567 Δ*lacZ*4787(::*rrnB*-3) *rph*-1 Δ(*rhaD*-*rhaB*)568 *hsdR*514], MG1655 (K-12, F^-^ λ^-^ *ilvG*^-^ *rfb*-50 *rph*-1), and Nissle 1917 were obtained from laboratory stocks. *E. cloni* 10G [K–12, F^-^ λ^-^ Δ(*ara leu*)7697 *araD*139 Δ*lacX*74 *galU galK* Φ80d *lacZ*ΔM15 *recA*1 *endA*1 *nupG rpsL* (Str^R^) Δ(*mrr*-*hsdRMS*-*mcrBC*) *tonA*] was obtained from Lucigen. 10G is a DH10B derivative and is suitable for maintaining large DNA constructs (Durfee et al., 2008). Bacterial strains IJ284 *Klebsiella* sp. 390 (O3:K11), IJ1668 K-12 hybrid; K1 capsule, and IJ612 *Salmonella typhimurium* LT2 were kindly provided by Ian Molineux. Virulence-plasmid-less *Yersinia pseudotuberculosis* IP2666 and YPIII were kindly provided by Joan Mecsas (Tufts University). *Pseudomonas aeruginosa* PAO1 was obtained from a laboratory stock. *E. coli* libraries, such as the ECOR group and DECA set, were sourced from the Thomas S. Whittam STEC Center (Michigan State University). *Pseudomonas putida* C1S (ATCC 23287) was obtained from ATCC. The pRS415 yeast centromere vector with *LEU2* marker (ATCC 87520) was obtained from a laboratory stock. Primers used in this study are listed in Table S5.

**Synthesis of codon-optimized 13a gene *17*.** The gene was synthesized by Gen9. The sequence is shown in Table S2.

**Culture conditions.** *S. cerevisiae* BY4741 was cultured in YPD [1% Bacto Yeast Extract (BD), 2% Bacto Peptone (BD), 2% dextrose (VWR)] at 30°C. *Y. pseudotuberculosis* strains and *P. putida* C1S were cultured in LB (BD) at 30°C. All other strains were cultured in LB at 37°C.

**Preparation of linearized pRS415 amplicon.** We linearized the pRS415 by using PCR amplification with specific primer sets (Table S5) and KAPA HiFi DNA Polymerase (Kapa Biosystems). For capturing phage genomes, we added sequences to the pRS415 vector that were homologous to the 5’ and 3’ termini of phages. To prevent the appearance of false-positive colonies, we excised and purified the pRS415 amplicon from an agarose gel after electrophoresis with QIAquick Gel Extraction Kit (Qiagen).

**Preparation of phage genomes.** Lysates were made by infecting 200 ml of logarithmically growing cells with the appropriate phage at a MOI of 0.1-0.01 and incubating the cultures until clearance. Cells were lysed and lysates were sterilized by adding 200 μl chloroform (Sigma). Lysates were centrifuged at 8,000 g for 5 min and then filtered through 0.22 μm filters (VWR) to remove cell debris. We added 216 μl of buffer L1 [20 mg/ml RNase A (Sigma), 6 mg/ml DNase I (NEB), 0.2 mg/ml BSA (NEB), 10 mM EDTA (Teknova), 100 mM Tris-HCl (VWR), 300 mM NaCl (VWR), pH 7.5] and incubated at 37°C for 1 h with gentle shaking. Then we added 30 ml of ice cold buffer L2 [30% polyethylene glycol (PEG) 6000 (Sigma), 3 M NaCl] and stored the samples overnight in 4°C. Samples were centrifuged at 10,000 g for 30 min at 4°C. Phage pellets were suspended in 9 ml buffer L3 (100 mM Tris-HCl, 100 mM NaCl, 25 mM EDTA, pH7.5). Then, we added 9 ml buffer L4 [4% SDS (VWR)] and incubated the samples at 70°C for 20 min. After cooling down on ice, 9 ml buffer L5 [2.55 M potassium acetate, pH4.8 (Teknova)] were added, and the samples were centrifuged at 10,000 g for 30 min at 4°C. Phage genomes in the supernatant were purified by using the Qiagen-tip 100 system (Qiagen) according to the manufacturer’s instructions.

**Preparation of PCR products for assembling phage genomes.** All PCR products were prepared with specific primer sets (Table S5) and KAPA HiFi DNA Polymerase. Five to ten 3.8-12.0 kbp PCR products including the YAC were used per reaction. Homology arms between the YAC and the phage genomes were added to the first and last phage genome fragments in order to decrease background recombination between the YAC and phage genomic DNA used as templates for PCR, and to eliminate the need for the time-consuming step of gel extraction for each PCR fragment.

**Preparation of yeast competent cells.** *S. cerevisiae* BY4741 was grown in 5 ml YPD at 30°C for 24 h. Overnight cultures were added into 50 ml YPD, and incubated at 30°C for 4 h. Cells were harvested by centrifugation at 3,000 g and washed with 25 ml water and then with 1 ml of 100 mM lithium acetate (LiAc) (Alfa Aesar), and suspended in 400 μl of 100 mM LiAc. Fifty microliters were used for each transformation.

**Yeast transformation.** All DNA samples and a linearized pRS415 were collected in a tube (0.5 - 4.0 μg for each DNA sample and 100 ng linearized pRS415 in 50 μl water), and mixed with the transformation mixture [50 μl yeast competent cell, 240 μl 50% PEG3350 (Sigma), 36 μl 1 M LiAc, 25 μl 2 mg/ml salmon sperm DNA (Sigma)]. The mixture was incubated at 30°C for 30 min, then at 42°C for 20 min or at 42°C for 45 min, centrifuged at 8,000 g for 15 sec, and suspended in 200 μl water. Transformants were selected on complete synthetic defined medium without leucine (SD-Leu) [0.67% YNB+Nitrogen (Sunrise Science Products), 0.069% CSM-Leu (Sunrise Science Products), 2% dextrose] agar plates at 30°C for 3 days.

**Extraction of captured phage genomes.** Individual yeast transformants were picked into SD-Leu liquid medium and incubated at 30°C for 24 h. DNA was extracted from these cells using the YeaStar Genomic DNA Kit (Zymo Research) or Yeast Genomic DNA Purification Kit (Amresco) according to the manufacturer’s instructions.

**Rebooting of phages.** The *E. coli* 10G strain was used as a host bacterium for the initial propagation of phages. To reboot T7 and T3 phages, 3 μl of extracted DNA were electroporated into 20 - 25 μl cells in a 2 mm gap electroporation cuvette (Molecular BioProducts) at 2,500 V, 25 μF, 200 Ω using a Gene Pulser Xcell (Bio-Rad). Cells were mixed with 3 ml LB soft agar (LB + 0.6% agar) warmed at 55°C, poured onto LB plate, and incubated for 4 h at 37°C. To reboot other phages, after electroporation, cells were incubated at 37°C for 1-2 h in 1 ml LB medium. Then, we added drops of chloroform to kill the cells and release phages. After centrifugation at 12,000 g for 1 min, supernatants were mixed with 300 μl overnight cultures of host bacteria for the phages and 3 ml LB soft agar, poured onto LB plate, and incubated for 4 - 24 h at 30 or 37°C.

**One-time phage propagation assays.** We used *E. coli* 10G from Lucigen as a one-time phage propagation host (Durfee et al., 2008). To validate the ability of the 10G strain as a one-time phage propagation plant, we electroporated 10 – 200 ng of purified phage genomes into the bacteria. After incubation for 1 - 2 h, which should be sufficient time for phages to have completed a full growth cycle, we added chloroform to kill the cells and release phages that may have failed to lyse cells. Then, supernatants were mixed with overnight cultures of each natural host bacteria for each phage in soft agar, poured onto agar plates, incubated for 4 - 24 h at 30 to 37°C, and analyzed for plaque formation.

**Determination of Plaque Forming Units (PFUs).** We mixed serially diluted phages in 0.95% saline, 300 μl overnight culture of host bacteria, and 3 ml LB soft agar, and poured the mixture onto LB plates. After 4 - 24 h incubation at 30 or 37°C, phage plaques were counted, and PFU/ml values were calculated.

**Plaque formation assays.** We mixed 300 μl bacterial overnight cultures and 3 ml LB soft agar, and poured the mixtures onto LB plate. After 5 min at room temperature (RT), 2.5 μl of 10-fold serially diluted phages in 0.95% saline were spotted onto LB soft agar and incubated at 30 or 37°C.

**Adsorption assay.** We mixed 100 μl of ∼10^8^ CFU/ml *E. coli* strains and T3 phage (MOI = 0.5), and incubated at RT for 10 min. Then, we added 700 μl of 0.95% saline and drops of chloroform to kill the cells and prevent the production of progeny phages. After centrifugation at 11,000 g for 1 min, supernatants were serially diluted and mixed with 300 μl of *E. coli* BL21 overnight cultures and 3 ml LB soft agar, and the mixtures were poured onto LB plates. After 4 h incubation at 37°C, phage plaques were counted, and adsorption efficiencies were calculated. Adsorption efficiency (%) = [1-(PFU of unadsorbed phage / original PFU in the BL21 and phage mixture)]×100

**Bacterial killing assays.** Overnight cultures of *Y. pseudotuberculosis* IP2666 and *Klebsiella* sp. 390 were diluted 1:200 into LB and grown to log-phase (≈10^8^ CFU/ml), i.e., for 5 h at 30°C and for 3 h at 37°C, respectively. Bacterial cultures were mixed with phage lysates (MOI ∼0.1) and incubated at 30 or 37°C. At each time point, bacteria were collected, washed twice with 0.95% saline, serially diluted, plated onto LB, and incubated at 30 or 37°C. Colonies were enumerated to calculate CFU/ml.

**Microbiome editing assays.** Overnight cultures of *E. coli* Nissle 1917, *Klebsiella* sp. 390, and *Y. pseudotuberculosis* IP2666 were diluted 1:200 into LB and grown to log-phase (∼10^8^ CFU/ml), i.e., for 3 h at 37°C, for 3 h at 37°C, and for 5 h at 30°C, respectively. Cultures were mixed and treated with phage lysates (MOI ∼0.1) and incubated at 30°C. At each time point, bacteria were collected, washed twice with 0.95% saline, serially diluted, plated onto LB, LB containing 25 μg/ml carbenicillin (VWR), and LB containing 1 μg/ml triclosan (VWR), and incubated at 30°C. Colonies were enumerated to calculate CFU/ml.

**Statistical analysis.** For all data points in all experiments, three samples were collected. The data are presented as the mean, and the error bars represent the SEM. In the “**Bacterial killing assays**” and the **“Microbiome editing assays”**, all CFU data were log_10_-transformed before analysis.

## AUTHOR CONTRIBUTIONS

H.A. and T.K.L. designed the study. H.A., S.L., and D.P.P. performed experiments. All authors analyzed the data and discussed results. H.A., S.L., and T.K.L. wrote the manuscript. H.A., S.L., and T.K.L. have filed a provisional application on this work.

## ACKNOWLEDGMENTS

Strains IJ284 *Klebsiella* sp. 390, IJ1668 K-12 hybrid; K1 capsule, IJ612 *S. typhimurium* LT2, K1E, K1F, K1-5, SP6, and K11 were kindly provided by Ian Molineux (The University of Texas at Austin). LUZ19 was kindly provided by Rob Lavigne (KU Leuven). *Yersinia pseudotuberculosis* IP2666 and YPIII were kindly provided by Joan Mecsas (Tufts University). We thank Oliver Purcell and Jennifer Henry for critical reading of the manuscript. This work was supported by grants from the Defense Threat Reduction Agency (HDTRA1-14-1-0007), the National Institutes of Health (1DP2OD008435, 1P50GM098792, 1R01EB017755), and the U. S. Army Research Laboratory and the U. S. Army Research Office through the Institute for Soldier Nanotechnologies, under contract number W911NF-13-D-0001. H.A. was supported by fellowships from the Japan Society for the Promotion of Science, and the Naito Foundation. D.P.P. was supported by the Portuguese Foundation for Science and Technology through the grant SFRH/BD/76440/2011.

## Supplemental Information

**Figure S1.**
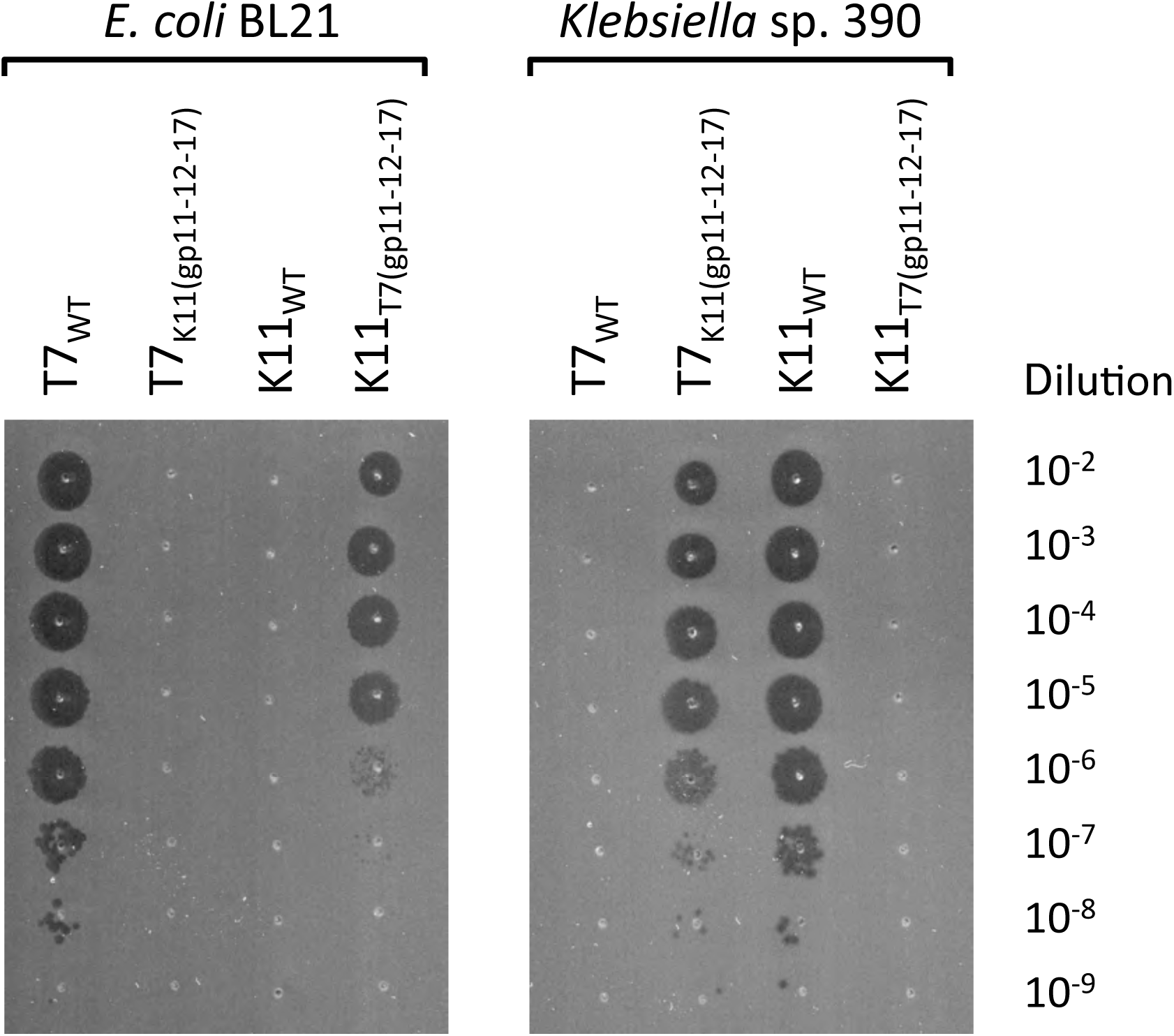
Plaque formation assays with T7_K11(gp11-12-17)_ and K11_T7(gp11-12-17)_, related to Figure 5. To confirm correct EOPs of T7_K11(gp11-12-17)_ and K11_T7(gp11-12-17)_ phages, 2.5 μL of 10- fold serially diluted phages were spotted onto bacterial lawns and incubated at 37°C. K11_T7(gp11-12-17)_ adopted the host range of T7_WT_ while T7_K11(gp11-12-17)_ adopted the host range of K11, thus demonstrating that tail component swapping can lead to acquisition of novel host ranges.

**Figure S2.**
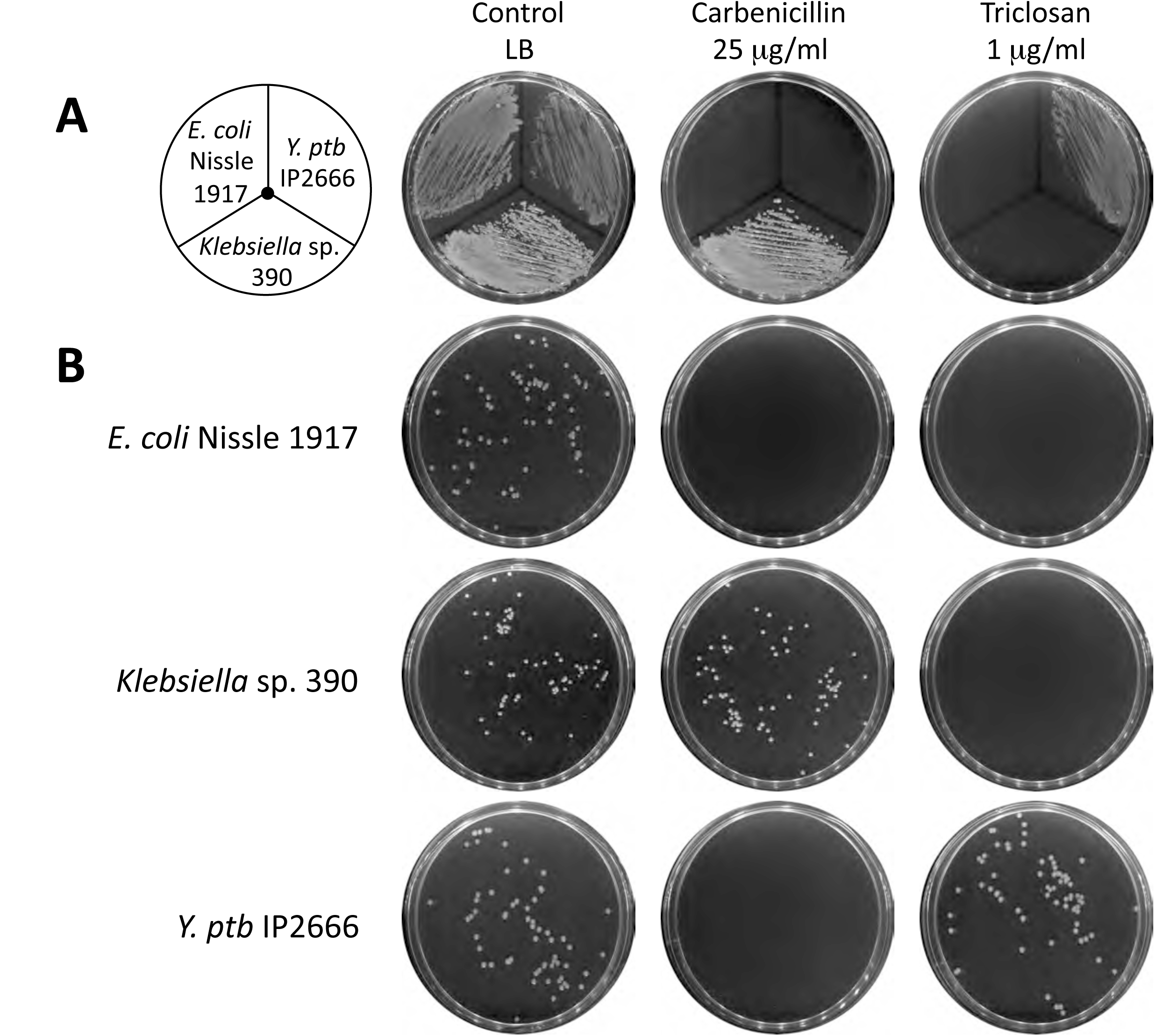
Antimicrobial susceptibilities of *E. coli* Nissle 1917, *Klebsiella* sp. 390, and *Y. pseudotuberculosis* IP2666, related to Figure 6. (**A**) Five microliters of each overnight cultures (>10^9^ CFU/ml) for each bacteria were streaked on LB plates with or without antibiotics. Plates were incubated at 30°C for 24 h. *Klebsiella* sp. 390 was naturally resistant to 25 mg/ml carbenicillin and *Y. ptb* IP2666 was naturally resistant to 1 mg/ml triclosan, while *E. coli* Nissle 1917 was sensitive to both. (**B**) Diluted log-phase cultures were plated onto LB with or without antibiotics. After incubation at 30°C for 18-24 h, colonies were enumerated.

**Table S1.** One-time phage propagation assay, related to Figure 1.

| Phage | Plaque formation on *E. coli* 10G | Propagation in *E. coli* 10G | Host bacteria |
| --- | --- | --- | --- |
| T7 | Yes | Yes | *E. coli* BL21 |
| T3 | Yes | Yes | *E. coli* BL21 |
| K1E | No | Yes | *E. coli* IJ1668 |
| K1F | No | Yes | *E. coli* IJ1668 |
| K1-5 | No | Yes | *E. coli* IJ1668 |
| SP6 | No | Yes | *S. typhimurium* IJ612 |
| LUZ19 | No | Yes | *P. aeruginosa* PAO1 |
| gh-1 | No | Yes | *P. putida* C1S |
| K11 | No | Yes | *Klebsiella* sp. 390 |

**Table S2.**
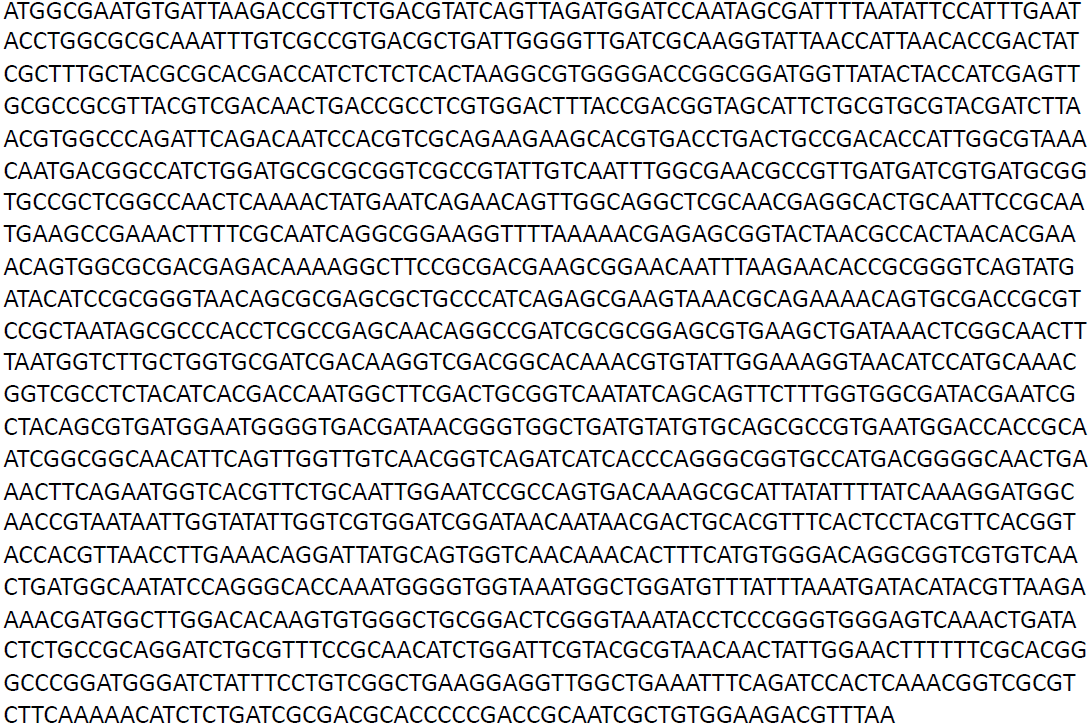
Sequence of codon-optimized 13a gene *17*, related to Figure 2E.

**Table S3.**
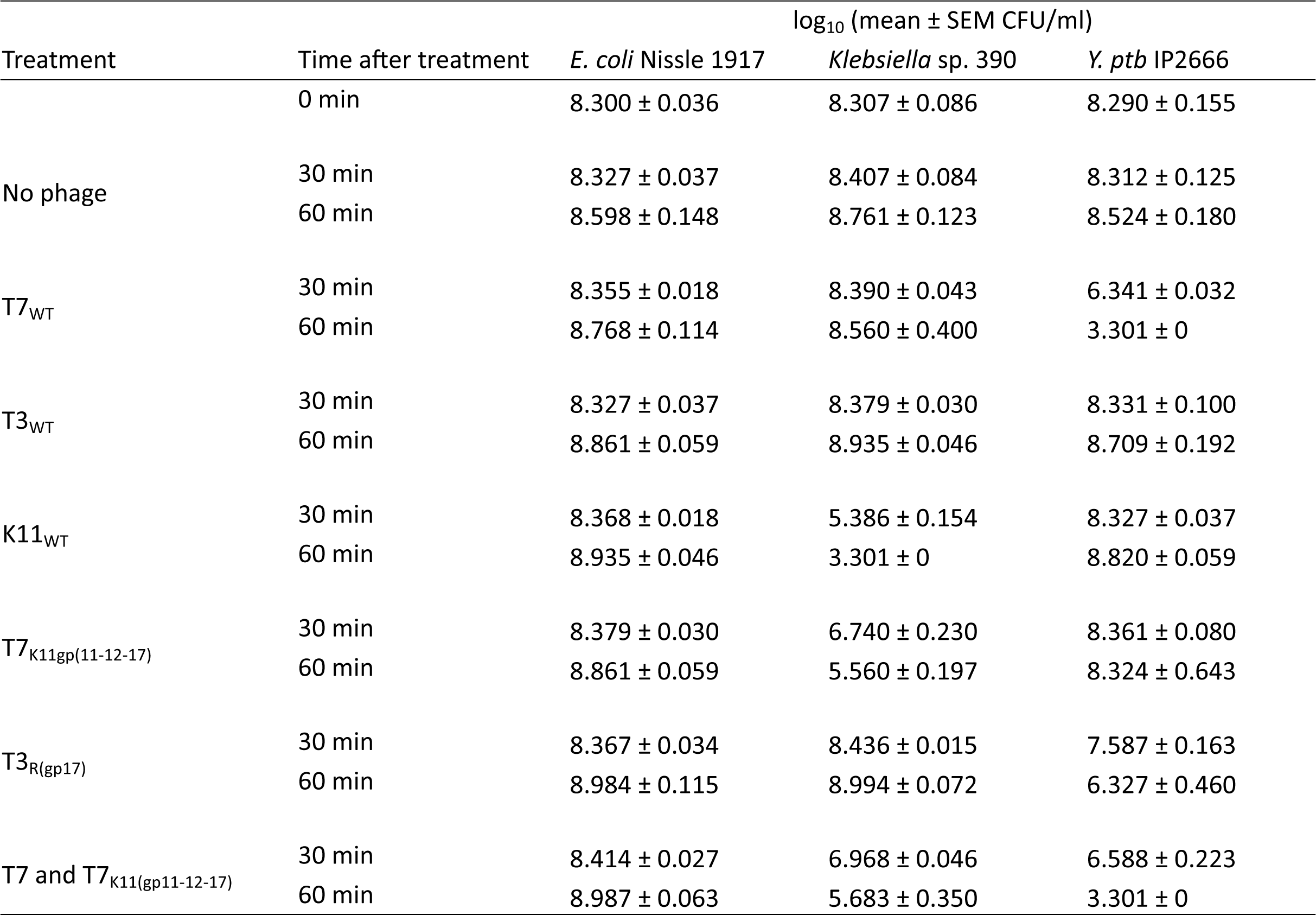
Microbiome editing assay, related to Figure 6.

**Table S4.** Synthetic phages created in this study, related to Experimental Procedures.

| Phage | Genotype | Description |
| --- | --- | --- |
| T7<sub>WT</sub> | T7 wild-type | Synthesized from PCR products |
| T3<sub>WT</sub> | T3 wild-type | Synthesized from PCR products |
| K1E<sub>WT</sub> | K1E wild-type | Synthesized from PCR products |
| K1F<sub>WT</sub> | K1F wild-type | Synthesized from PCR products |
| K1-5<sub>WT</sub> | K1-5 wild-type | Synthesized from PCR products |
| SP6<sub>WT</sub> | SP6 wild-type | Synthesized from PCR products |
| gh-1<sub>WT</sub> | gh-1 wild-type | Synthesized from PCR products |
| K11<sub>WT</sub> | K11 wild-type | Synthesized from PCR products |
| T7<sub>T3(C-gp17)</sub> | T7<sub>WT</sub> gene 17 (1-447)-T3 gene 17 (448-1677) | T7 with T7-T3 hybrid tail fiber |
| T7<sub>T3(gp17)</sub> | T7<sub>WT</sub> Δgene 17, T3 gene 17 | T7 with T3 tail fiber |
| T7<sub>13a(C-gp17)</sub> | T7<sub>WT</sub> gene 17 (1-450)-13a gene 17 (451-1677) | T7 with T7-13a hybrid tail fiber |
| T7<sub>13a(gp17)</sub> | T7<sub>WT</sub> Δgene 17, 13a gene 17 | T7 with 13a tail fiber |
| T7<sub>K11(gp11-12-17)</sub> | T7<sub>WT</sub> Δgene (11 12 17), K11 gene (11 12 17) | T7 with K11 tail |
| T3<sub>T7(C-gp17)</sub> | T3<sub>WT</sub> gene 17(1-447)-T7 gene 17(448-1662) | T3 with T3-T7 hybrid tail fiber |
| T3<sub>T7(gp17)</sub> | T3<sub>WT</sub> Δgene 17, T7 gene 17 | T3 with T7 tail fiber |
| T3<sub>R(gp17)</sub> | T3<sub>WT</sub> Δgene 17, R gene 17 | T3 with R tail fiber |
| K11<sub>T7(gp11-12-17)</sub> | K11<sub>WT</sub> Δgene (11 12 17), T7 gene (11-12 17) | K11 with T7 tail |

**Table S5.**
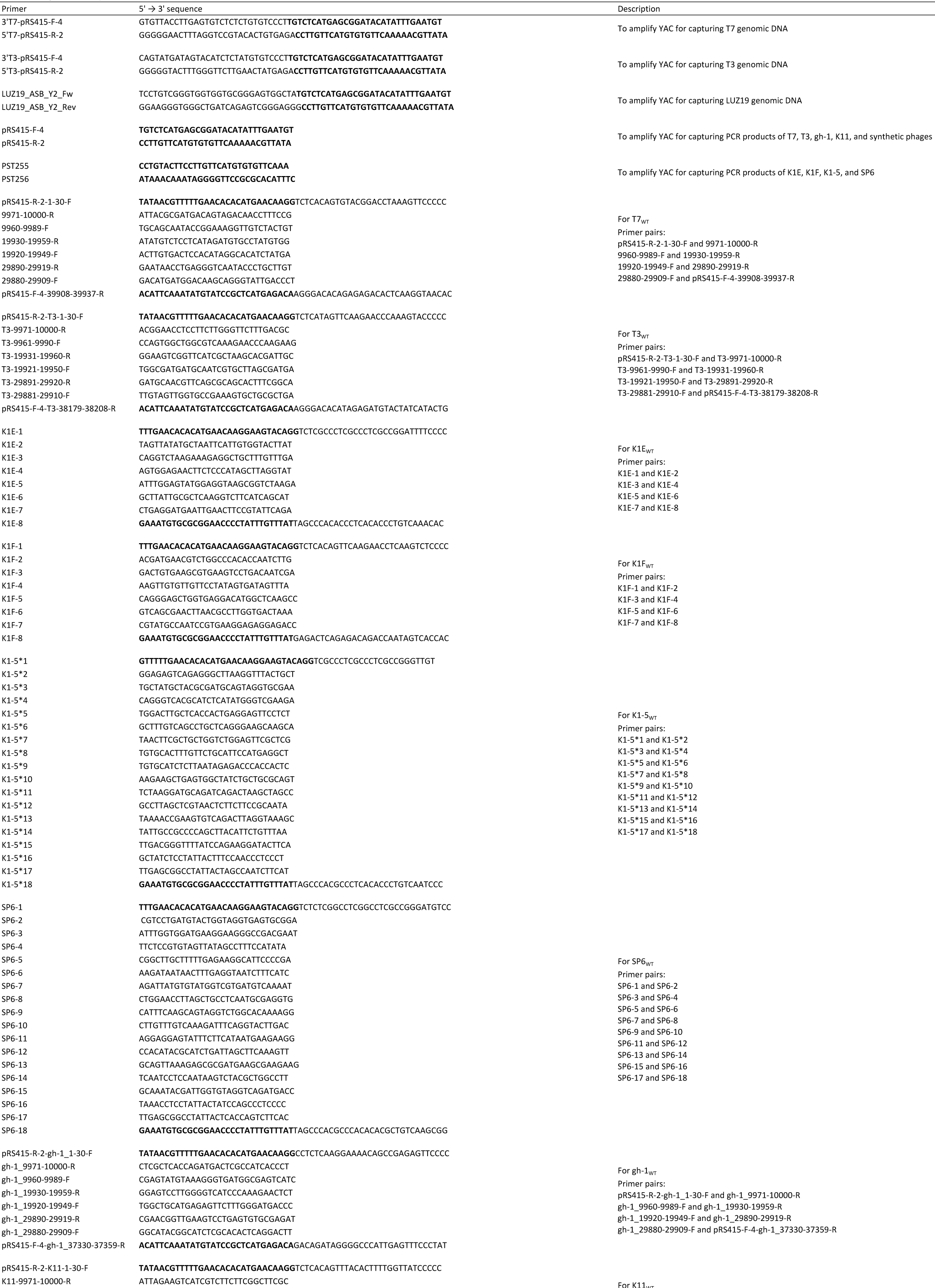
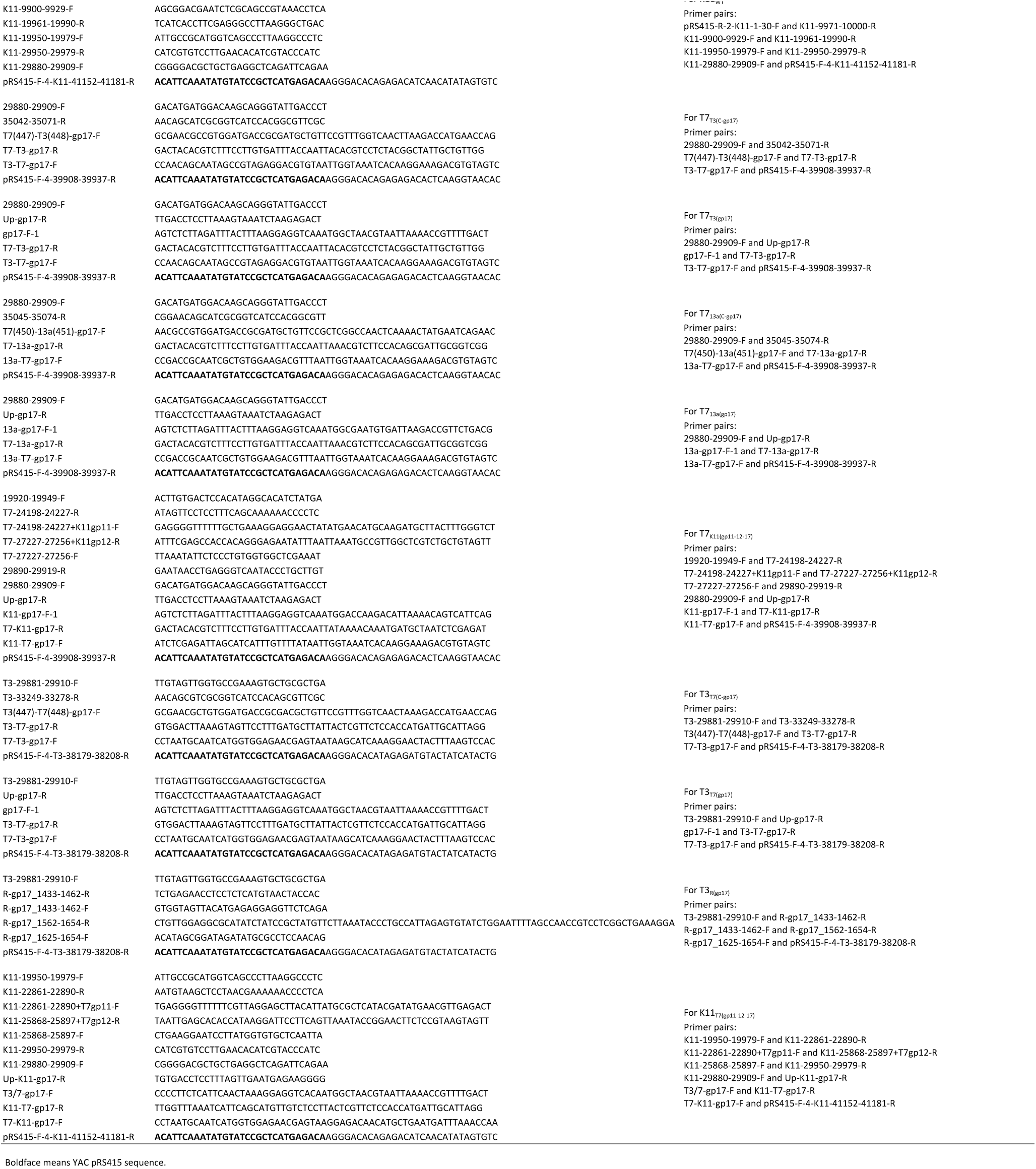
Oligonucleotide primers used in this study, related to Experimental Procedures.

## REFERENCES

Beier, H., Golomb, M., and Chamberlin, M. (1977). Isolation of recombinants between T7 and T3 bacteriophages and their use in vitro transcriptional mapping. Journal of virology 21, 753–765.

Bessler, W., Freund-Molbert, E., Knufermann, H., Rduolph, C., Thurow, H., and Stirm, S. (1973). A bacteriophage-induced depolymerase active on *Klebsiella* K11 capsular polysaccharide. Virology 56, 134–151.

Bikard, D., Euler, C.W., Jiang, W., Nussenzweig, P.M., Goldberg, G.W., Duportet, X., Fischetti, V.A., and Marraffini, L.A. (2014). Exploiting CRISPR-Cas nucleases to produce sequence-specific antimicrobials. Nature biotechnology 32, 1146–1150.

Brussow, H. (2012). What is needed for phage therapy to become a reality in Western medicine? Virology 434, 138–142.

Carlton, R.M. (1999). Phage therapy: past history and future prospects. Archivum immunologiae et therapiae experimentalis 47, 267–274.

Citorik, R.J., Mimee, M., and Lu, T.K. (2014). Sequence-specific antimicrobials using efficiently delivered RNA-guided nucleases. Nature biotechnology 32, 1141–1145.

Cuervo, A., Pulido-Cid, M., Chagoyen, M., Arranz, R., Gonzalez-Garcia, V.A., Garcia-Doval, C., Caston, J.R., Valpuesta, J.M., van Raaij, M.J., Martin-Benito, J., et al. (2013). Structural characterization of the bacteriophage T7 tail machinery. The Journal of biological chemistry 288, 26290–26299.

d’Herelle, F. (1931). Bacteriophage as a Treatment in Acute Medical and Surgical Infections. Bulletin of the New York Academy of Medicine 7, 329–348.

Demerec, M., and Fano, U. (1945). Bacteriophage-Resistant Mutants in *Escherichia Coli*. Genetics 30, 119–136.

Dietz, A., Weisser, H.J., Kossel, H., and Hausmann, R. (1990). The gene for *Klebsiella* bacteriophage K11 RNA polymerase: sequence and comparison with the homologous genes of phages T7, T3, and SP6. Molecular & general genetics : MGG 221, 283–286.

Dunn, J.J., and Studier, F.W. (1983). Complete nucleotide sequence of bacteriophage T7 DNA and the locations of T7 genetic elements. Journal of molecular biology 166, 477–535.

Durfee, T., Nelson, R., Baldwin, S., Plunkett, G., 3rd, Burland, V., Mau, B., Petrosino, J.F., Qin, X., Muzny, D.M., Ayele, M., et al. (2008). The complete genome sequence of *Escherichia coli* DH10B: insights into the biology of a laboratory workhorse. Journal of bacteriology 190, 2597–2606.

Fischbach, M.A., and Walsh, C.T. (2009). Antibiotics for emerging pathogens. Science 325, 1089–1093.

Garcia, E., Elliott, J.M., Ramanculov, E., Chain, P.S., Chu, M.C., and Molineux, I.J. (2003). The genome sequence of *Yersinia pestis* bacteriophage phiA1122 reveals an intimate history with the coliphage T3 and T7 genomes. Journal of bacteriology 185, 5248–5262.

Gibson, D.G. (2012). Oligonucleotide assembly in yeast to produce synthetic DNA fragments. Methods in molecular biology 852, 11–21.

Gibson, D.G., Benders, G.A., Axelrod, K.C., Zaveri, J., Algire, M.A., Moodie, M., Montague, M.G., Venter, J.C., Smith, H.O., and Hutchison, C.A., 3rd (2008). One-step assembly in yeast of 25 overlapping DNA fragments to form a complete synthetic *Mycoplasma genitalium* genome. Proceedings of the National Academy of Sciences of the United States of America 105, 20404–20409.

Gibson, D.G., Young, L., Chuang, R.Y., Venter, J.C., Hutchison, C.A., 3rd, and Smith, H.O. (2009). Enzymatic assembly of DNA molecules up to several hundred kilobases. Nature methods 6, 343–345.

Goldberg, G.W., Jiang, W., Bikard, D., and Marraffini, L.A. (2014). Conditional tolerance of temperate phages via transcription-dependent CRISPR-Cas targeting. Nature 514, 633–637.

Grice, E.A., and Segre, J.A. (2012). The human microbiome: our second genome. Annual review of genomics and human genetics 13, 151–170.

Hendrix, R.W. (2003). Bacteriophage genomics. Current opinion in microbiology 6, 506–511.

Jaschke, P.R., Lieberman, E.K., Rodriguez, J., Sierra, A., and Endy, D. (2012). A fully decompressed synthetic bacteriophage oX174 genome assembled and archived in yeast. Virology 434, 278–284.

Kiro, R., Shitrit, D., and Qimron, U. (2014). Efficient engineering of a bacteriophage genome using the type I-E CRISPR-Cas system. RNA biology 11, 42–44.

Lin, A., Jimenez, J., Derr, J., Vera, P., Manapat, M.L., Esvelt, K.M., Villanueva, L., Liu, D.R., and Chen, I.A. (2011). Inhibition of bacterial conjugation by phage M13 and its protein g3p: quantitative analysis and model. PLoS one 6, e19991.

Lin, T.Y., Lo, Y.H., Tseng, P.W., Chang, S.F., Lin, Y.T., and Chen, T.S. (2012). A T3 and T7 recombinant phage acquires efficient adsorption and a broader host range. PLoS one 7, e30954.

Lu, T.K., Koeris, M.S., Chevalier, B.S., Holder, J.W., Mckenzie, G.J., and Brownell, D.R. (2013). Recombinant phage and methods. Patent WO 2013049121 A2.

Ma, H., Kunes, S., Schatz, P.J., and Botstein, D. (1987). Plasmid construction by homologous recombination in yeast. Gene 58, 201–216.

Molineux, I.J. (2006). The T7 Group. In The Bacteriophages, R. Calendar, ed. (New York: Oxford Univ. Press), pp. 277–301.

Pajunen, M.I., Elizondo, M.R., Skurnik, M., Kieleczawa, J., and Molineux, I.J. (2002). Complete nucleotide sequence and likely recombinatorial origin of bacteriophage T3. Journal of molecular biology 319, 1115–1132.

Pouillot, F., Blois, H., and Iris, F. (2010). Genetically engineered virulent phage banks in the detection and control of emergent pathogenic bacteria. Biosecurity and bioterrorism : biodefense strategy, practice, and science 8, 155–169.

Qimron, U., Tabor, S., and CC., R. (2010). New details about bacteriophage T7-host interactions. In Microbe magazine.

Rashid, M.H., Revazishvili, T., Dean, T., Butani, A., Verratti, K., Bishop-Lilly, K.A., Sozhamannan, S., Sulakvelidze, A., and Rajanna, C. (2012). A *Yersinia pestis*-specific, lytic phage preparation significantly reduces viable *Y. pestis* on various hard surfaces experimentally contaminated with the bacterium. Bacteriophage 2, 168–177.

Shin, J., Jardine, P., and Noireaux, V. (2012). Genome replication, synthesis, and assembly of the bacteriophage T7 in a single cell-free reaction. ACS synthetic biology 1, 408–413.

Steven, A.C., Trus, B.L., Maizel, J.V., Unser, M., Parry, D.A., Wall, J.S., Hainfeld, J.F., and Studier, F.W. (1988). Molecular substructure of a viral receptor-recognition protein. The gp17 tail-fiber of bacteriophage T7. Journal of molecular biology 200, 351–365.

Sulakvelidze, A., Alavidze, Z., and Morris, J.G.Jr., (2001). Bacteriophage therapy. Antimicrob Agents Chemother 45, 649–659.

Tetart, F., Desplats, C., and Krisch, H.M. (1998). Genome plasticity in the distal tail fiber locus of the T-even bacteriophage: recombination between conserved motifs swaps adhesin specificity. Journal of molecular biology 282, 543–556.

Trojet, S.N., Caumont-Sarcos, A., Perrody, E., Comeau, A.M., and Krisch, H.M. (2011). The gp38 adhesins of the T4 superfamily: a complex modular determinant of the phage’s host specificity. Genome biology and evolution 3, 674–686.

Yaung, S.J., Esvelt, K.M., and Church, G.M. (2014). CRISPR/Cas9-mediated phage resistance is not impeded by the DNA modifications of phage T4. PLoS one 9, e98811.

Yoichi, M., Abe, M., Miyanaga, K., Unno, H., and Tanji, Y. (2005). Alteration of tail fiber protein gp38 enables T2 phage to infect *Escherichia coli* O157:H7. Journal of biotechnology 115, 101–107.

